# Reconstituting spore cortex peptidoglycan biosynthesis reveals a deacetylase that catalyzes transamidation

**DOI:** 10.1101/2023.03.12.532300

**Authors:** Micaela J. Tobin, Stephen Y. Cho, William Profy, Tessa M. Ryan, Donna H. Le, Crystal Lin, Elaine Z. Yip, Jack L. Dorsey, Blake R. Levy, Jillian D. Rhodes, Michael A. Welsh

**Affiliations:** Chemistry Department, Hamilton College, 198 College Hill Rd. Clinton, NY 13323

## Abstract

Some bacteria survive in nutrient-poor environments and resist killing by antimicrobials by forming spores. The cortex layer of the peptidoglycan cell wall that surrounds mature spores contains a unique modification, muramic-δ-lactam, that is essential for spore germination and outgrowth. Two proteins, the amidase CwlD and the deacetylase PdaA, are required for muramic-δ-lactam synthesis in cells, but their combined ability to generate muramic-δ-lactam has not been directly demonstrated. Here we report an in vitro reconstitution of cortex peptidoglycan biosynthesis, and we show that CwlD and PdaA together are sufficient for muramic-δ-lactam formation. Our method enables characterization of the individual reaction steps, and we show for the first time that PdaA has transamidase activity, catalyzing both the deacetylation of *N*-acetylmuramic acid and cyclization of the product to form muramic-δ-lactam. This activity is unique among peptidoglycan deacetylases and is notable because it may involve the direct ligation of a carboxylic acid with a primary amine. Our reconstitution products are nearly identical to the cortex peptidoglycan found in spores, and we expect that they will be useful substrates for future studies of enzymes that act on the spore cortex.

## MAIN TEXT

In response to nutrient starvation, many species of Gram-positive bacteria form spores that can germinate, reinitiating vegetative growth when permissive conditions are established.^1^ Spores are metabolically dormant bacterial cells with a modified external cell envelope and a dehydrated cytoplasm.^2^ These features make the spore highly resistant to killing by both physical and chemical means, including antibiotic treatment. Indeed, spore formation is the basis for transmission, chronic infection, and antibiotic evasion by human pathogens such as *Clostridioides difficile*.^3^ Characterization of the molecular steps involved in sporulation and germination is therefore of interest because these pathways may be promising targets for new spore-specific antimicrobials.

Bacteria are surrounded by an essential cell wall that is composed of peptidoglycan, a carbohydrate polymer of alternating *N*-acetylglucosamine (GlcNAc) and *N*-acetylmuramic acid (MurNAc) residues (Figure 1).^4, 5^ In vegetative cells, each MurNAc is appended with a pentapeptide stem that serves as the site of covalent crosslinking between polymer strands. Mature spores contain two distinct layers of peptidoglycan.^2^ A thin, inner layer, called the germ cell wall, is similar in composition to that of vegetative cells and is retained during spore germination and outgrowth. In contrast, a thick, outer layer of peptidoglycan, the cortex, is fully degraded during spore germination and contains a distinct chemical modification. Roughly half of the MurNAc residues are deacetylated, their stem peptides removed, and the resulting muramic acid (MurN) residue cyclized to give muramic-δ-lactam (Figure 1).^6–9^ This modification serves as a recognition motif that recruits hydrolases to the cortex during germination, targeting it for destruction.^10^ Thus, germination is greatly impaired in spores without muramic-δ-lactam.^11^

**Figure 1.**
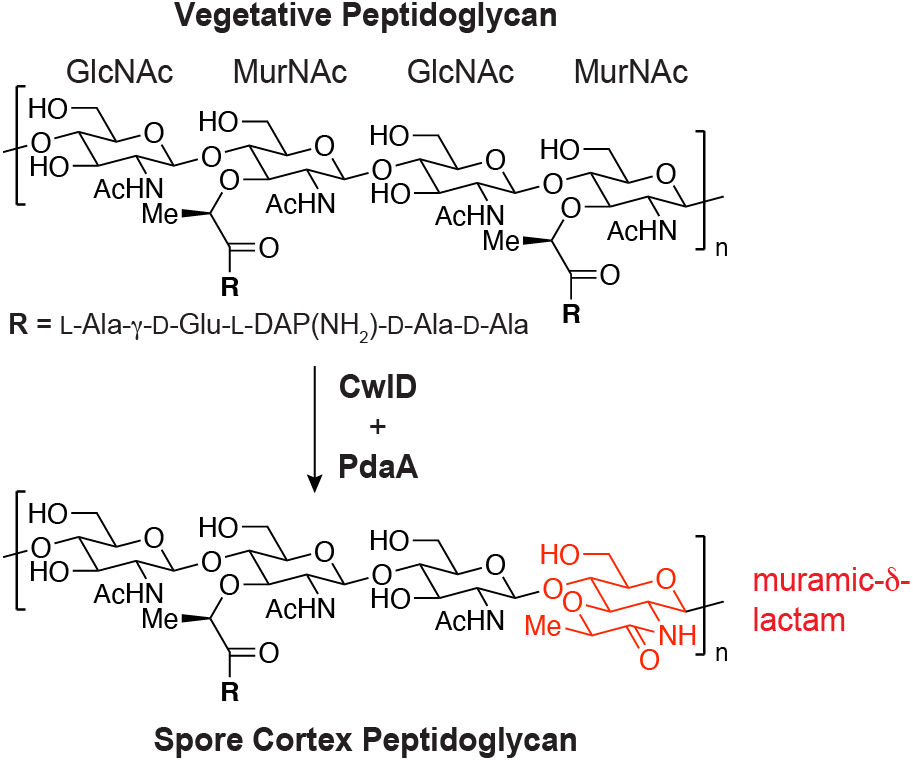
Structure of spore cortex peptidoglycan. In mature spores, up to 50% of MurNAc residues are converted to muramic-δ-lactam. The remaining stem peptides (R) are frequently truncated or part of covalent crosslinks between polymer strands.

Although muramic-δ-lactam was first identified in spore peptidoglycan decades ago,^6^ its biosynthesis has lacked full characterization. At least two enzymes are required for muramic-δ-lactam production in the model organism *Bacillus subtilis*, the L-alanine amidase CwlD and the MurNAc deacetylase PdaA (Figure 1).^12, 13^ In prior work, heterologous expression of the *B. subtilis* proteins in the periplasm of *Escherichia coli* revealed that muramic-δ-lactam was formed only when both CwlD and PdaA were present.^14^ Expression of CwlD alone produced peptide-cleaved MurNAc while expression of PdaA alone resulted in no detectable change to the peptidoglycan.^14^ These findings have led to a model for muramic-δ-lactam synthesis whereby CwlD acts first to remove the stem peptide followed by amine deacetylation and lactam cyclization by PdaA, but this model has not been directly verified in vitro. To our knowledge, CwlD has not been reconstituted, and biochemical work on PdaA to date has confirmed only its deacetylase activity; lactam ring formation was not observed.^15^ Which protein catalyzes the cyclization, or if it is uncatalyzed, remains an open question.

Reconstitution of enzymes that act on peptidoglycan is challenging because many of these proteins will only use large peptidoglycan oligomers as substrates. Producing these substrates by chemical synthesis is nontrivial, but recent advances in obtaining the peptidoglycan precursor Lipid II have made it possible to prepare structurally defined peptidoglycan substrates in vitro.^16^ We reasoned that providing CwlD and PdaA with a native peptidoglycan substrate may allow us to reconstitute muramic-δ-lactam synthesis and characterize their activities. CwlD and PdaA variants from two sporulating model organisms, *B. subtilis* and *Clostridioides difficile*, were overexpressed in *E. coli* and purified to homogeneity (Figure S1). To generate a suitable substrate for the enzymes, we extracted Lipid II from *B. subtilis* and polymerized it into linear peptidoglycan strands using SgtB, a monofunctional glycosyltransferase from *Staphylococcus aureus*.^17^ We then incubated the individual CwlD or PdaA proteins with the linear polymer and analyzed the reaction products using a liquid chromatography-mass spectrometry (LC-MS) assay. In this analysis, polymeric products are digested into smaller fragments with the muramidase mutanolysin and reduced with sodium borohydride (NaBH4) to enable LC-MS detection. Mutanolysin digestion of unmodified peptidoglycan gives disaccharide products (Figure 2a, product A, Figure S2), but mutanolysin cleaves MurNAc-β(1→4)-GlcNAc linkages only when the MurNAc bears a pentapeptide. Stem peptide removal by an amidase therefore produces an altered cleavage pattern, giving tetrasaccharide or longer fragments (Figures 2a,b, product B, Figure S2). Reactions with *B. subtilis* PdaA, *Bs*PdaA, alone gave only starting material A (Figure 2c, i); we did not detect any modified disaccharide or tetrasaccharide products. This result is in good agreement with prior work suggesting that unmodified peptidoglycan is not a substrate for PdaA.^14, 15^ Reactions with only *B. subtilis* CwlD, *Bs*CwlD, produced B as the major product, confirming that this protein acts as an amidase (Figure 2c, ii). To determine if the CwlD product is a substrate for PdaA, we treated linear peptidoglycan with *Bs*CwlD for 1 h followed by addition of *Bs*PdaA. We observed the near-disappearance of B along with the appearance of three new products (Figure 2c, iii). The major product, C, contained MurN (Figure 2b, S2), indicating that PdaA could deacetylate MurNAc only once the stem peptide had been removed. Product D contained muramic-δ-lactam, but the peak was small. However, we also detected another product, E, with a mass consistent with reduction of muramic-δ-lactam by NaBH4 (Figure 2b, S2). While most amides do not react with NaBH_4_, the decalin-like lactam is sufficiently strained to be reduced.^6–8^ Product E is therefore derived from D and is evidence of muramic-δ-lactam formation. When we allowed the *Bs*PdaA reaction to run for 48 h, we observed E as the major product, occurring in yields of up to 75% (Fig 2c, iv). Analogous experiments conducted with the CwlD and PdaA variants from *C. difficile, Cd*CwlD and *Cd*PdaA1, produced similar results although the amidase activity of *Cd*CwlD was weak (Figure S3). This is presumably because our reactions did not contain GerS, an activator of *Cd*CwlD required for sporulation in this organism.^18, 19^ Nevertheless, *Cd*PdaA1 activity was robust, with all product B converted to C or lactam (Figure S3).

**Figure 2.**
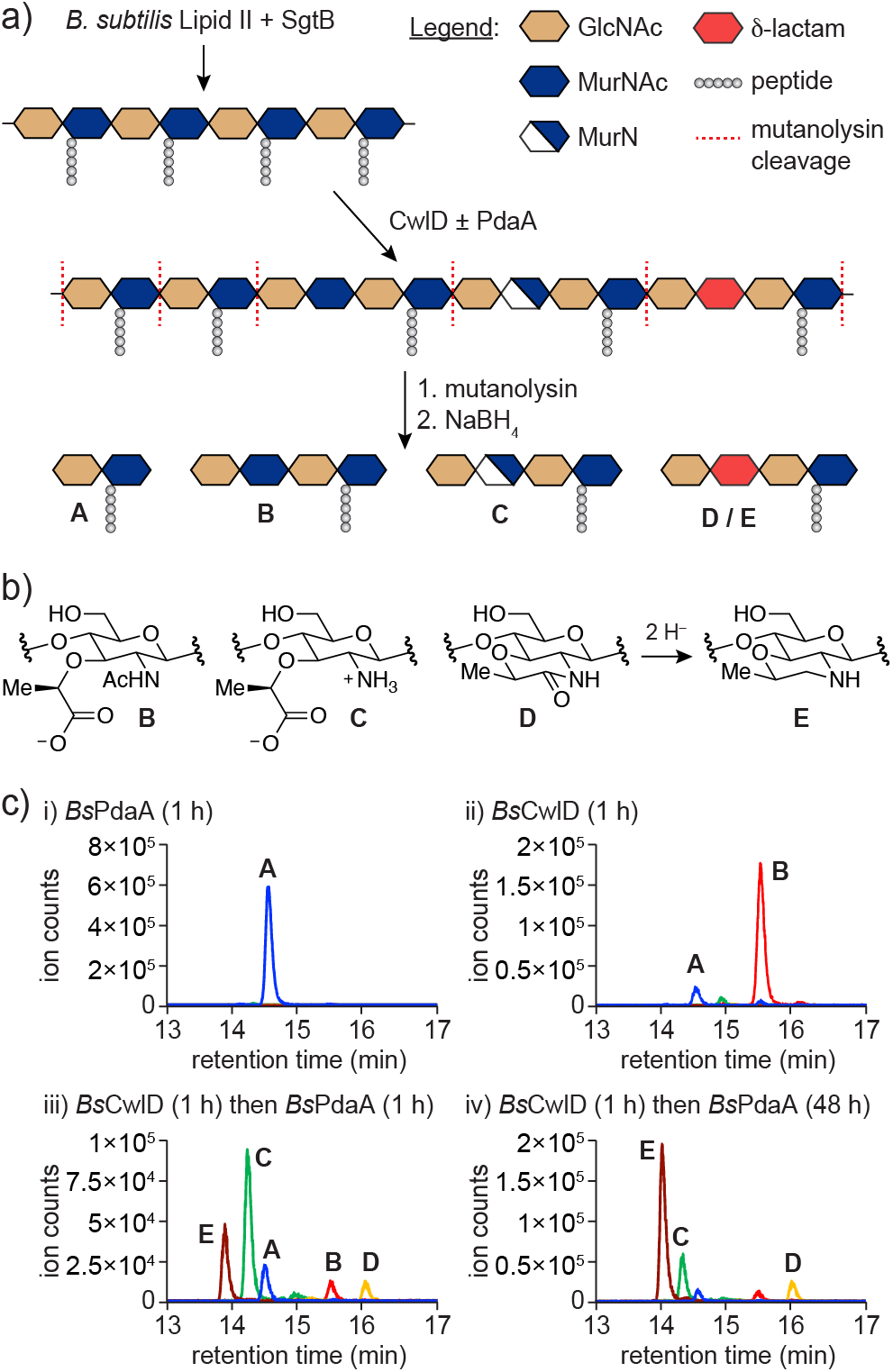
Reconstitution of muramic-δ-lactam synthesis. (a) Schematic of the LC-MS assay used to characterize the peptidoglycan products of CwlD and PdaA reactions. (b) Chemical structures of the modified MurN residues found in products B, C, D, and E. (c) LC-MS analysis of *Bs*CwlD and *Bs*PdaA reactions. Linear peptidoglycan was incubated with the indicated enzyme in pH 7.5 buffer. The data is representative of at least three independent experiments.

Together, these results show that CwlD and PdaA are sufficient for muramic-δ-lactam synthesis. They also confirm the prevailing model whereby the synthesis is ordered; CwlD acts first followed by PdaA. Importantly, the products produced on longer PdaA reaction times (24-48 h) are consistent with peptidoglycan where approximately 40% of MurNAc residues have been converted to muramic-δ-lactam (see Supporting Methods) and with the modified residues occurring every other monomer. This distribution is similar to what is observed in the cortex layer of spores where 30-50% of residues are muramic-δ-lactam, depending on the bacterium.^7–9^ We expect that the ability to generate defined cortex peptidoglycan substrates will prove useful for studying other enzymes that act on the cell wall during sporulation and germination. We note that in our initial reactions with PdaA, we found that the lactam synthesis was maximal when Ca^2+^ ions were present in the reaction buffer (Figure S4). The absence of these ions could explain why only deacetylase activity, not lactam cyclization, was observed previously.^15^ Because Ca^2+^ would be abundant in the intermembrane compartment where PdaA localizes,^20^ it is plausible that these ions are required to stabilize a conformation that permits lactam synthesis.

Having successfully reconstituted muramic-δ-lactam formation, we next sought to determine the relative rates of deacetylation and lactam cyclization under our reaction conditions. CwlD and PdaA are both metalloenzymes that contain a divalent cation cofactor, often Zn^2+^, used to promote the nucleophilicity of a water molecule (Figure S5).^18, 21^ Their reactions can therefore be quenched by addition of excess chelator such as EDTA (Figure S6).^22, 23^ To assess activity over time, we treated linear peptidoglycan with *Bs*CwlD to generate polymer enriched in peptide-cleaved product B and incubated the resulting material with *Bs*PdaA or *Cd*PdaA1. Reaction aliquots were EDTA-quenched and analyzed via our LC-MS method (Figure 3a). Both PdaA variants catalyzed relatively rapid deacetylation as evidenced by a burst of product C within the first 5-10 min along with a corresponding decrease in B (Figure 3b,c). Muramic-δ-lactam began to appear within the first 30 min and accumulated gradually over several hours, corresponding with a decrease in product C. These results suggest that MurN, the product of PdaA deacetylase activity, could be a reaction intermediate that is subsequently cyclized to muramic-δ-lactam.

**Figure 3.**
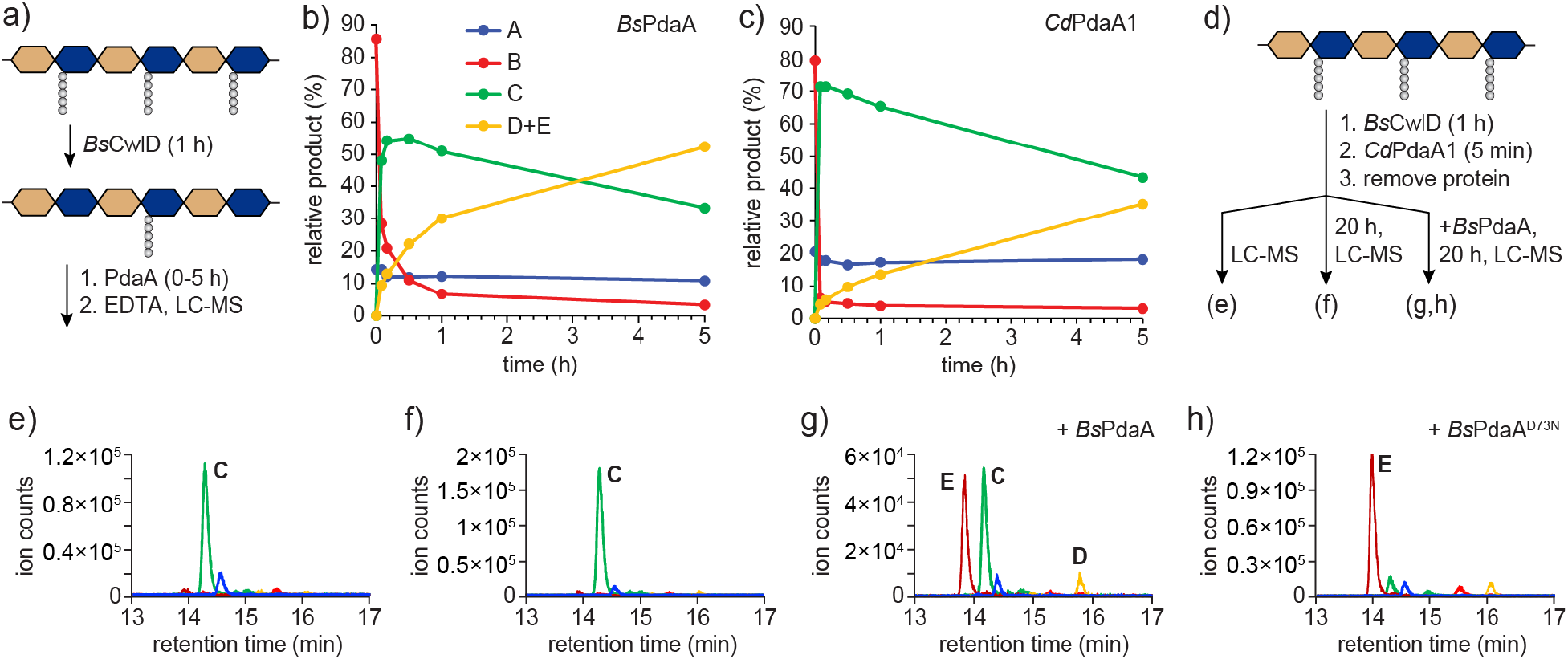
PdaA catalyzes cyclization of MurN to muramic-δ-lactam. (a) Schematic of the workflow for PdaA timecourse analysis. *Bs*PdaA (b) and *Cd*PdaA1 (c) reactions were monitored by LC-MS over 5 h. (d) Schematic of the MurN cyclization assay. Peptidoglycan enriched in MurN was generated by successive treatment of linear polymer with *Bs*CwlD and *Cd*PdaA1 followed by removal of the proteins on Ni-NTA resin. Eluted polymer products were characterized by LC-MS immediately (e), after 20 h at room temperature (f), or after re-addition of *Bs*PdaA or *Bs*PdaA^D73N^ for 20 h (g,h). All reactions were conducted in pH 7.5 buffer. Data is representative of at least three independent experiments.

We next sought to determine if the lactam cyclization was catalyzed by CwlD or PdaA. We noted in the timecourse analysis that *Cd*PdaA1 generated polymer greatly enriched in MurN, product C, at early timepoints (Figure 3c). We envisioned that if the proteins could be removed following production of MurN, we would have generated a substrate we could use to assay the cyclization step of the reaction. To do this, we prepared linear peptidoglycan and added *Bs*CwlD for 1 h followed by *Cd*PdaA1 for 5 min. The reaction mixture was then passed over a plug of Ni-NTA resin to remove the His6-tagged proteins (Figure 3d, S7). LC-MS analysis of the eluted peptidoglycan showed polymer enriched in product C (Figure 3e). When we allowed the eluted product to sit at room temperature for 20 h, we observed no change in the product distribution (Figure 3f); thus, uncatalyzed cyclization of MurN is slow under our reaction conditions. Readdition of *Bs*CwlD to the eluted material did not result in lactam synthesis (Figure S8); however, re-addition of *Bs*PdaA gave good conversion of MurN to muramic-δ-lactam (Figure 3g). Analogous experiments with *Cd*PdaA1 resulted in almost complete cyclization of MurN (Figure S8). From these results, we conclude that the lactam cyclization is catalyzed by PdaA. Our data are therefore consistent with a model for muramic-δ-lactam synthesis shown in Figure 4a.

**Figure 4.**
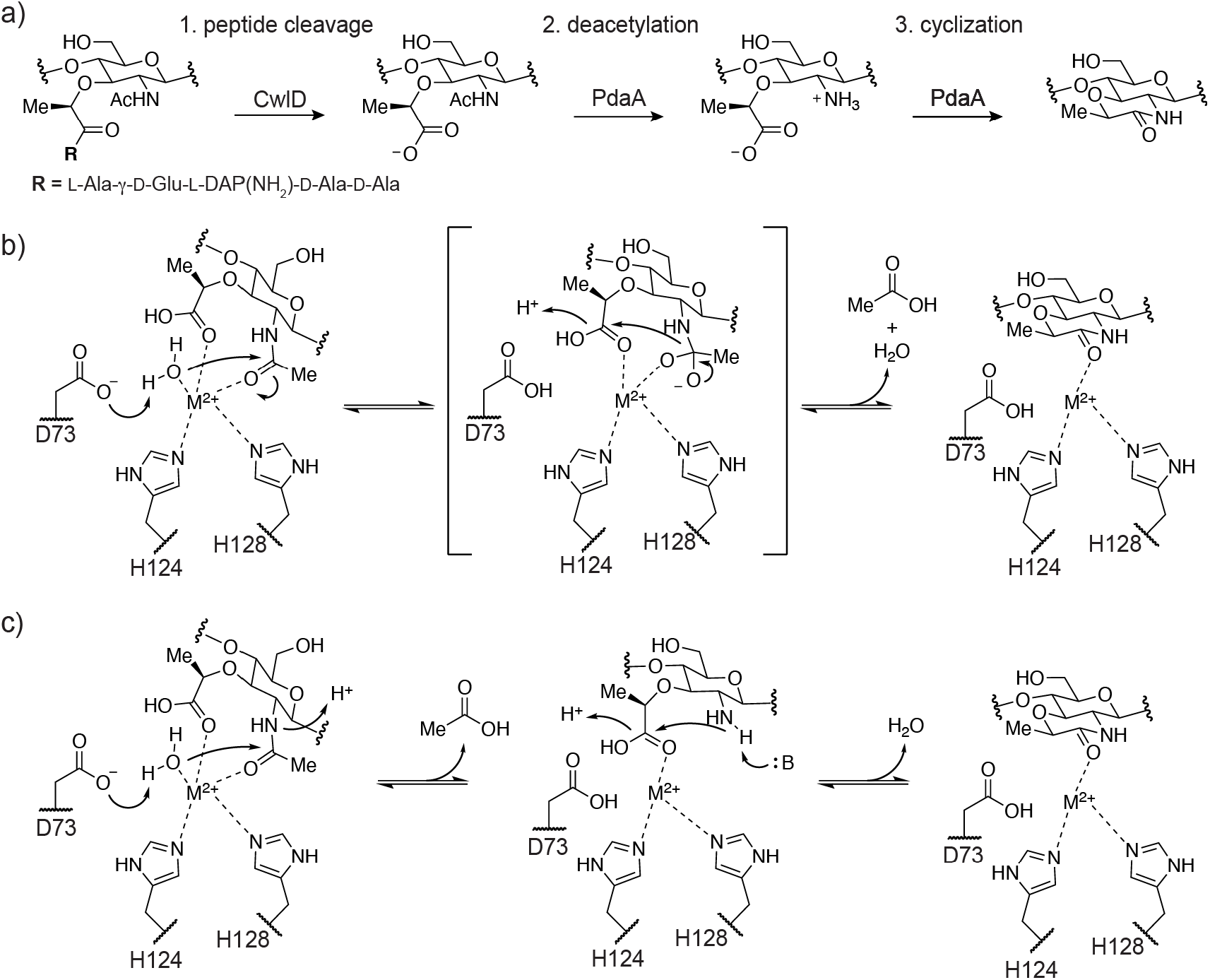
The biosynthetic pathway for muramic-δ-lactam. (a) The amidase CwlD catalyzes stem peptide cleavage. This is followed by deacetylation and cyclization by PdaA. (b) A possible transamidation mechanism for *Bs*PdaA whereby lactam is formed after a single substrate binding step. (c) An alternate mechanism involving two substrate-binding steps that is consistent with our data. Deacetylation is followed by substrate release then re-binding and cyclization. For brevity, not all steps are shown.

The reaction catalyzed by PdaA, a net transamidation, is remarkable in several respects. PdaA is part of a larger family of enzymes bearing a NodB homology domain (InterPro IPR002509) and classified as carbohydrate esterase 4 (CE4) in the CAZy database.^24, 25^ All other characterized members of this enzyme family act solely as deacetylases or esterases. Therefore, the transamidase activity exhibited by PdaA is unique. Notably though, lactam cyclization does not depend on deacetylase activity. D73 of *Bs*PdaA acts as a catalytic base in the deacetylation reaction.^22, 23^ A D73N mutation abolished deacetylase activity (Figure S9), but the mutant protein was able to catalyze cyclization of a previously deacetylated substrate (Figure 3h). How PdaA accomplishes lactam cyclization merits further investigation. Possible reaction mechanisms for transamidation are shown in Figure 4b,c. Although we cannot rule out other options at this stage, the mechanism shown in Figure 4c is consistent with our observation that the cyclization can proceed directly through MurN (Figure 3g,h). This was surprising because direct ligation of a carboxylate and amine would be energetically unfavorable, yet muramic-δ-lactam formation does not require input of chemical energy, such as ATP, to generate an intermediate activated for acyl-substitution. How then is PdaA able to catalyze direct amide bond formation? Binding of MurN into a reactive conformation, combined with desolvation and charge neutralization of the reacting groups, could be sufficient to accelerate the reaction at room temperature. The MurN residue is a privileged substrate for such a reaction given that the carboxyl and amine groups are tethered and the product is a six-membered ring. We are unaware of enzymes aside from PdaA that catalyze direct amide bond formation, and we are currently conducting further investigation into this unusual protein.

## ASSOCIATED CONTENT

### Supporting Information

The Supporting Information is available free of charge. Contents include detailed experimental methods, bacterial strains, plasmids, and supporting figures and tables.

### UniProt Accession Codes

*Bs*CwlD: P50864, *Bs*PdaA: O34928, *Cd*CwlD: Q18CJ4, *Cd*PdaA1: Q18BV2

## AUTHOR INFORMATION

### Author Contributions

M.A.W. led the study. All authors participated in data collection. M.A.W. prepared figures and wrote the paper. ‡These authors contributed equally.

## ACKNOWLEDGMENT

We thank Hamilton College for providing startup funds that supported this work. M.A.W. is grateful to Suzanne Walker for hosting his sabbatical leave. We thank Bailey Schultz, Eric Snow, and Suzanne Walker for critical reading of the manuscript. We thank Dan Kahne and Ryan Martinie for helpful discussions.

