## Supplemental Information for "Reconstituting spore cortex peptidoglycan biosynthesis reveals a deacetylase that catalyzes transamidation"

Chemistry Department, Hamilton College, Clinton, New York 13323

**METHODS**

**Materials.** Unless indicated otherwise, all reagents and chemicals were purchased from Sigma-Aldrich and used without further purification. Oligonucleotide PCR primers were purchased from Integrated DNA Technologies. *B. subtilis* Lipid II was extracted and purified as reported.[1] *S. aureus* SgtB was purified as previously described.[2] Mini-PROTEAN TGX gels (4-20%, Bio-Rad) were used for SDS-PAGE.

**Bacterial Culture.** The bacterial strains and plasmids used in this study are listed in Table S1. *E. coli* strains were grown at 37 °C with shaking in LB broth (Beckton Dickinson) or on agarized plates. Antibiotics were used at the following concentrations: ampicillin, 100 μg/mL; carbenicillin, 100 μg/mL; kanamycin, 50 μg/mL.

**Cloning of *B. subtilis* CwlD.** Plasmid pET28b(+) was linearized by PCR using primers oMW26 and oMW27 (Table S2). The *cwlD[N27-E237]* gene, lacking the predicted N-terminal signal peptide, was amplified from *B. subtilis* PY79 genomic DNA using the primers oMW29 and oMW30. The DNA products were gel purified and joined by isothermal assembly using In-Fusion Master Mix (Takara Bio) to produce plasmid pMW1008, which expresses *B. subtilis* CwlD[N27-E237] with an N-terminal $His_6$-tag. The *cwlD[N27-E237]* insert was confirmed by Sanger sequencing.

**Cloning of *B. subtilis* PdaA and PdaA$^{D73N}$.** Plasmid pET24b was linearized by PCR using primers oMW47 and oMW48 (Table S2). The *pdaA[V24-L263]* gene, lacking the predicted N-terminal signal peptide, was amplified from *B. subtilis* PY79 genomic DNA using the primers oMW66 and oMW67. The DNA products were gel purified and joined by isothermal assembly using Gibson Assembly Master Mix (New England Biolabs) to produce plasmid pMW1070, which expresses *B. subtilis* PdaA[V24-L263] with a C-terminal $His_6$-tag. The *pdaA[V24-L263]* insert was confirmed by Sanger sequencing.

To generate the D73N mutant, pMW1070 was amplified using primers oMW124 and oMW125, which encode the desired mutation. The linear product was circularized using KLD reaction mix (New England Biolabs) to produce plasmid pMW1159-6, which expresses *B. subtilis* PdaA[V24-L263] (D73N) with a C-terminal $His_6$-tag.

**Cloning of *C. difficile* CwlD.** Plasmid pET28b(+) was linearized by PCR using primers oMW26 and oMW27 (Table S2). The *cwlD[S29-S234]* gene, lacking the predicted N-terminal signal peptide, was amplified from *C. difficile* 630 genomic DNA (ATCC) using the primers oMW34 and oMW35. The DNA products were gel purified and joined by isothermal assembly using In-Fusion Master Mix (Takara Bio) to produce plasmid pMW1015, which expresses *C. difficile* CwlD[S29-S234] with an N-terminal $His_6$-tag. The *cwlD[S29-S234]* insert was confirmed by Sanger sequencing.

**Cloning of *C. difficile* PdaA.** Plasmid pET28b(+) was linearized by PCR using primers oMW26 and oMW27 (Table S2). The *pdaA[S29-K247]* gene, lacking the predicted N-terminal signal peptide, was amplified from *C. difficile* 630 genomic DNA (ATCC) using the primers oMW36 and oMW37. The DNA products were gel purified and joined by isothermal assembly using In-Fusion Master Mix (Takara Bio) to produce plasmid pMW1016, which expresses *C.*

*difficile* PdaA[S29-K247] with an N-terminal $His_6$-tag. The *pdaA[S29-K247]* insert was confirmed by Sanger sequencing.

**Expression and purification of CwlD and PdaA – general protocol used for all proteins.** *E. coli* C43(DE3) containing the appropriate plasmid was grown in 500 mL LB broth supplemented with kanamycin at 37 ºC with shaking until the OD600 was 0.5-0.6. The culture was cooled to 16 ºC before inducing protein expression with 500 µM IPTG and shaking for 16 h. Cells were harvested by centrifugation (4,200 × g, 15 min, 4 ºC) and resuspended in 20 mL lysis buffer (20 mM Tris pH 7.5, 400 mM NaCl). The cell suspension was supplemented with DNase (0.5 mg/mL) and phenylmethanesulfonyl fluoride (1 mM) and the cells lysed by passage through a cell disruptor (EmulsiFlex-C3, Avestin) at ≥ 10,000 psi. Cell debris was pelleted by centrifugation (20,000 × *g*, 30 min, 4 ºC). The resulting supernatant was supplemented with 40 mM imidazole and then rocked with 0.5 mL Ni-NTA resin (Qiagen) for 45 min at 4 ºC. The resin was collected in a column by gravity flow and then washed twice with 5 mL wash buffer (20 mM Tris pH 7.5, 400 mM NaCl, 40 mM imidazole). The protein was eluted in 10 mL elution buffer (20 mM Tris pH 7.5, 400 mM NaCl, 200 mM imidazole) and concentrated by centrifugal filtration. The protein was then further purified by fast protein liquid chromatography (FPLC, AKTA Pure, Cytiva) on a Superdex 200 Increase 10/300 column (Cytiva) in a running buffer consisting of 20 mM Tris pH 7.5, 400 mM NaCl. Pooled elution fractions were concentrated by centrifugal filtration and the protein absorbance measured at 280 nm. The predicted extinction coefficient of the protein (via ProtParam[3]) was used to estimate concentration. Protein was diluted to 200 µM in running buffer with 10% glycerol (v/v), aliquoted, and stored at -80 ºC.

**CwlD and PdaA reactions – general conditions.** *B. subtilis* Lipid II was polymerized with SgtB, a monofunctional peptidoglycan glycosyltransferase from *Staphylococcus aureus*. Pooled polymerization reactions of up to 1 mL total volume were assembled under the following conditions: 50 mM HEPES, pH 7.5, 2 mM $CaCl_2$, 20 µM Lipid II, 0.2 µM SgtB, 10% DMSO (v/v). Reactions were incubated at room temperature for 30 min. CwlD and/or PdaA were added at 2 µM and the reactions incubated static at room temperature. Reactions were quenched by addition of EDTA at a final concentration of 10 mM and the mixture immediately frozen at -20 ºC.

**Digestion and LC-MS analysis of reaction products.** Peptidoglycan products of CwlD/PdaA reactions were digested with mutanolysin to enable LC-MS analysis. To 50 µL aliquots of peptidoglycan reaction products, 8 U of mutanolysin was added and the reaction incubated at 37 ºC for 2 h. Aqueous sodium borohydride (10 mg/mL, 50 µL) was added and the reaction incubated for 30 min at room temperature. The solution pH was adjusted to ~4 by addition of 20% phosphoric acid (approximately 5 µL) and the reactions lyophilized to dryness. The residue was dissolved in 25 µL of water and analyzed by LC-MS.

LC-MS was conducted using an Agilent Technologies 1200 series HPLC in line with an Agilent 6520 Q-TOF mass spectrometer using electrospray ionization and operating in positive ion mode. Products were separated on a Waters Symmetry Shield RP18 column (5 µm, 3.9 × 150 mm) with matching column guard using the following method: 0.4 mL/min solvent A (water/0.1% formic acid) for 5 min followed by a linear gradient of 0 to 40% solvent

B (acetonitrile/0.1% formic acid) over 25 min. The peptidoglycan fragments were observed to elute between 13-17 min. Masses of the predicted $[M+H]^+$ and $[M+2H]^{+2}$ ions for the expected muropeptide fragments were extracted from the resulting total ion chromatograms to generate the extracted ion chromatograms displayed in the text and supplemental figures. Mass spectrometry data was analyzed using Agilent MassHunter Workstation Qualitative Analysis software version B.06.00.

**Estimation of total conversion of MurNAc to muramic-δ-lactam.** Co-incubation of linear peptidoglycan with *Bs*CwlD and *Bs*PdaA for 24-48 h yielded polymeric products where approximately 40% of MurNAc residues were converted to lactam. Product A contains one unmodified MurNAc residue. Products B, C, D, and E each contain one modified and one unmodified MurNAc residue. The percent of MurNAc residues converted to muramic-δ-lactam was therefore calculated according to the formula below, where each letter indicates the integrated area of the respective peak.

$$\%\ lactam = \left(\frac{D + E}{A + 2B + 2C + 2D + 2E}\right) \times 100$$

**MS/MS of digested peptidoglycan.** A pooled reaction (500 μL) to generate linear *B. subtilis* peptidoglycan was prepared under the general reaction conditions described above. A 50 μL aliquot was removed and frozen at –20 ºC (the "peak A" sample). *Bs*CwlD (2 μM) was added and the reaction incubated at room temperature for 1 h. A 100 μL aliquot was removed, EDTA added at 10 mM, and the sample frozen at –20 ºC (the "peak B" sample). *Cd*PdaA1 (2 μM) was added to the pooled mixture and the reaction incubated at room temperature for 5 min. A 100 μL aliquot was removed, EDTA added at 10 mM, and the sample frozen at –20 ºC (the "peak C" sample). The remaining reaction mixture was incubated at room temperature for 20 h then split into two 100 μL aliquots. Mutanolysin (8 U) was added to all samples and the reactions incubated at 37 ºC for 2 h. To generate the "peak D" sample, 100 μL of 2 mg/mL aqueous sodium borohydride was added to an aliquot of 20 h reaction products and the reaction incubated at room temperature for 30 min. All other samples were reduced similarly with 10 mg/mL sodium borohydride with the second aliquot of 20 h reaction products representing the "peak E" sample. The pH of all reactions was adjusted to ~4 by addition of 20% phosphoric acid and the samples lyophilized to dryness. The residue was dissolved in 25 μL of water. The muropeptide reaction products were analyzed using the same LC-MS method and instrumentation as above. MS/MS was conducted in positive ion mode. For the peak A sample, the $[M+H]^+$ ion was targeted for fragmentation. The $[M+2H]^{+2}$ ion was targeted for all other samples. Ions were fragmented by collision-induced dissociation (CID) with the collision energy calculated according to the following formula: CID energy = (slope · m/z) / (100 + offset), where slope = 3.1 and offset = 2.

**Optimization of buffer conditions for PdaA.** For reactions investigating the effect of metal cations on PdaA activity (Figure S4), pooled reactions were assembled as described in the generic reaction conditions, above, but in buffer without $CaCl_2$. *Bs*CwlD (2 μM) was added and the reaction incubated 4 h at room temperature. EDTA was added 2 mM to quench the CwlD reaction and the mixture split into 50 μL aliquots. Frozen stocks of PdaA (200 μM) were combined 1:1 with 50 mM HEPES pH 7.5, 4 mM EDTA buffer and the solution incubated at 0 ºC for 15 min to strip the protein of metal cations. The PdaA solution was then added to the CwlD products to give a final PdaA concentration of 2 μM. Metal salt solutions ($MgCl_2$, $CaCl_2$, $MnCl_2$, $ZnSO_4$) were then added at 4 mM to initiate the reaction and the mixtures incubated at room temperature for 20 h. EDTA (10 mM) was added to quench and the sample frozen at -20 ºC. Mutanolysin digestion and LC-MS analysis were conducted as described above.

**Timecourse analysis of PdaA reactions.** To assess PdaA activity over time, a pooled Lipid II polymerization reaction was assembled as described in the generic reaction conditions, above. *Bs*CwlD (2 μM) was added and the reaction incubated 1 h at room temperature. A PdaA variant (2 μM) was then added and the reaction incubated static at room temperature. At timepoints, a 50 μL aliquot was removed, EDTA (10 mM) added, and the sample frozen at -20 ºC. Mutanolysin digestion and LC-MS analysis were conducted as described above. Relative product amounts were calculated by integrating the product peaks and dividing by the total peak area.

**Lactam Cyclization Assay.** To generate linear peptidoglycan substrates enriched in muramic acid residues (product C), pooled reactions were assembled as described in the generic reaction conditions, above. Linear *B. subtilis* peptidoglycan was treated with *Bs*CwlD (2 μM) for 1 h at room temperature. *Bs*PdaA or *Cd*PdaA1 (2 μM) was then added for 10 min and the mixture immediately passed over a 100 μL plug of settled Ni-NTA resin (Qiagen) by gravity flow. The eluate was passed over the resin a second time to maximize protein binding. The eluate was then split into 50 μL aliquots and treated as indicated in the main text. CwlD or PdaA variants were re-added at 2 μM and the resulting mixture incubated at room temperature for 20 h. All reactions were quenched by addition of 10 mM EDTA and stored at -20 ºC until needed. Mutanolysin digestion and LC-MS analysis were conducted as described above.

**Western Blotting.** Proteins separated by SDS-PAGE were transferred to a PVDF membrane. The membrane was blocked in tris-buffered saline with 5% dry milk (w/v) and 0.05% tween-20 (v/v) for 1 h at room temperature. $His_6$-tagged proteins were detected by incubation with THE™ His Tag Antibody [HRP] (Genscript, 1:5000 in blocking buffer) for 1 h at room temperature. The membrane was washed with tris-buffered saline, 0.05% tween-20. His-tagged proteins were detected by incubating the membrane in ECL reagent (ThermoFisher) and measuring chemiluminescence using an Azure 500 imager (Azure Biosystems).

### SUPPLEMENTAL FIGURES AND TABLES

**Table S1: Bacterial strains and plasmids**

| Strain or plasmid | Description[a] | Reference |
|---|---|---|
| *E. coli* | | |
| C43(DE3) | BL21(DE3) derivative for protein expression | 4 |
| *Plasmids* | | |
| pET24b | IPTG-inducible protein expression vector; $Kan^R$ | Novagen |
| pET28b(+) | IPTG-inducible protein expression vector; $Kan^R$ | Novagen |
| pMW1008 | *B. subtilis* $His_6$-CwlD[N27-E237] expression vector, $Kan^R$ | This study |
| pMW1015 | *C. difficile* $His_6$-CwlD[S29-S234] expression vector, $Kan^R$ | This study |
| pMW1016 | *C. difficile* $His_6$-PdaA[S29-K242] expression vector, $Kan^R$ | This study |
| pMW1070 | *B. subtilis* PdaA[V24-L263]-$His_6$ expression vector, $Kan^R$ | This study |
| pMW1159-6 | *B. subtilis* PdaA(D73N)-$His_6$ expression vector, $Kan^R$ | This study |
| pMgt1[†] | *S. aureus* SgtB-$His_6$ expression vector; $Amp^R$ | 5 |

[a] Abbreviations: $Amp^R$, ampicillin resistance; $Kan^R$, kanamycin resistance
[†] SgtB is also known as Mgt1.

**Table S2: Oligonucleotide PCR primers**

| primer | Sequence (5'-3') |
|---|---|
| oMW26 | CTCGAGCACCACCACCACC |
| oMW27 | CATATGGCTGCCGCGCGG |
| oMW29 | GTGGTGGTGCTCGAGTTACTCCGGAGGGTCTCCT |
| oMW30 | CGCGGCAGCCATATGAATAACGACTCTTGGAAGCCGTG |
| oMW34 | GTGGTGGTGCTCGAGTTAACTTAAATATTTTTGTATTCCT |
| oMW35 | CGCGGCAGCCATATGTCTGAAGATGTTATCAAGTATATGC |
| oMW36 | CGCGGCAGCCATATGTCATTAGATAAAACAAAAACT |
| oMW37 | GTGGTGGTGCTCGAGTTATTTTTTATTATTTAAATAATCA |
| oMW47 | CTCGAGCACCACCACCAC |
| oMW48 | CATATGTATATCTCCTTCTTAAAGTTAAACAAAATTATTTC |
| oMW66 | AAGAAGGAGATATACATATGGTGCCGAATGAGCCGATTAATTG |
| oMW67 | TGGTGGTGGTGGTGCTCGAGCAAAGACGGCAGCCTCATTTC |
| oMW124 | TAAATCGTTTTTTCCTTTGTATTC |
| oMW125 | CTTAACGTTTAATAACGGATATGAAAATG |

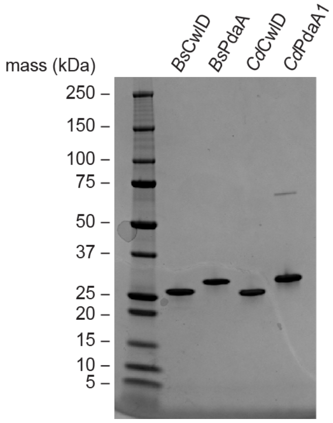


**Figure S1.** Coomassie-stained SDS-PAGE of purified proteins. Each lane contains 2 μg of protein.

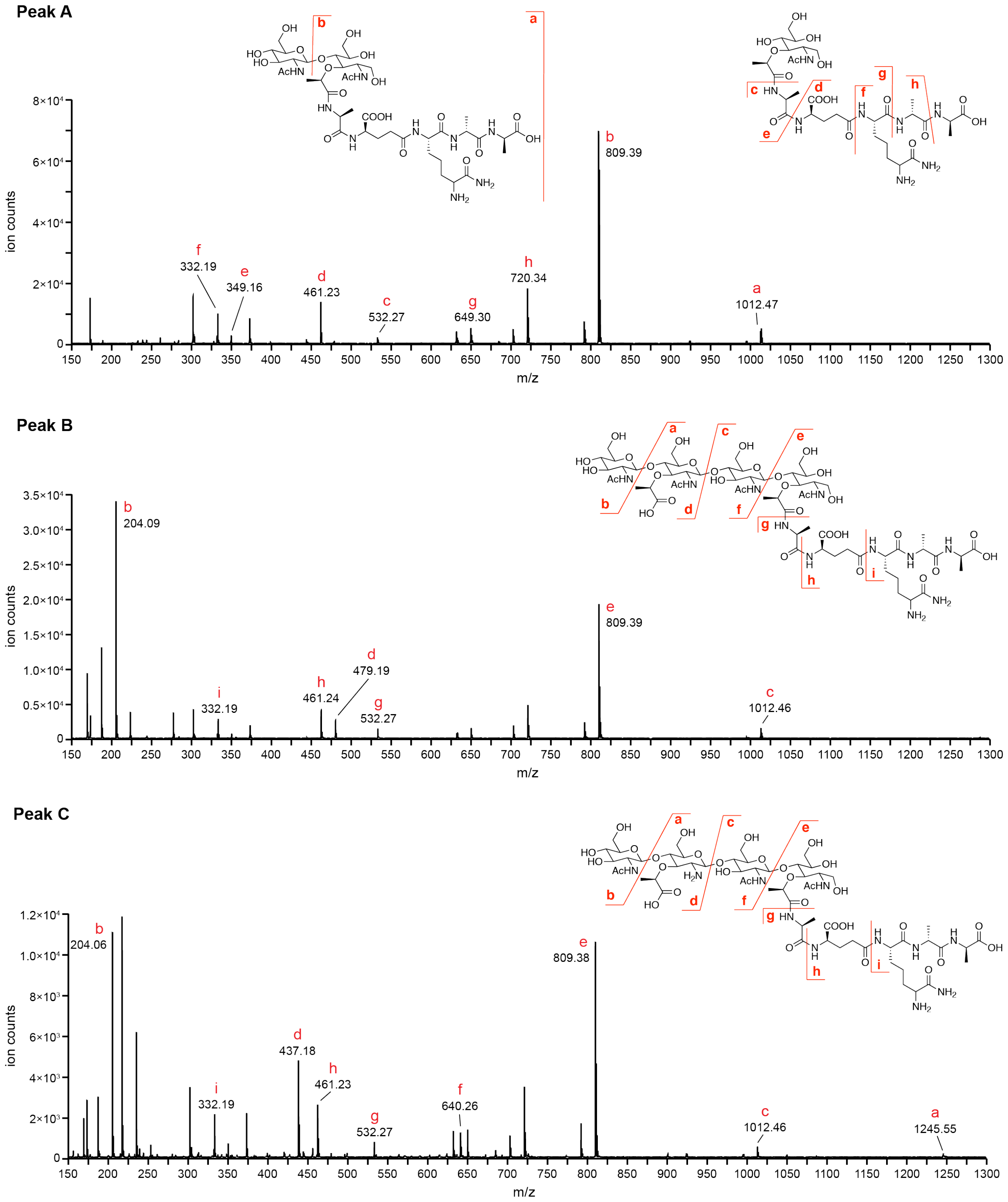


**Figure S2.** Targeted MS/MS of the peptidoglycan products formed by CwlD and PdaA. *B. subtilis* Lipid II was polymerized with SgtB and the resulting polymer treated with *Bs*CwlD and/or *Bs*PdaA as described in the Methods above. Products A, B, C, D, and E are defined as in the main text.

**Peak D** – muramic-δ-lactam fragment

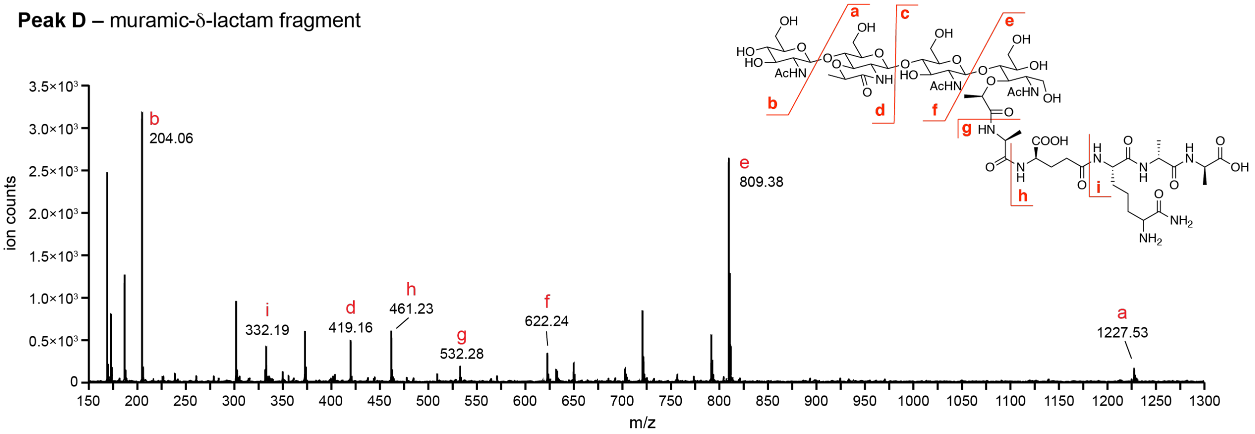


**Peak E** – reduced muramic-δ-lactam fragment

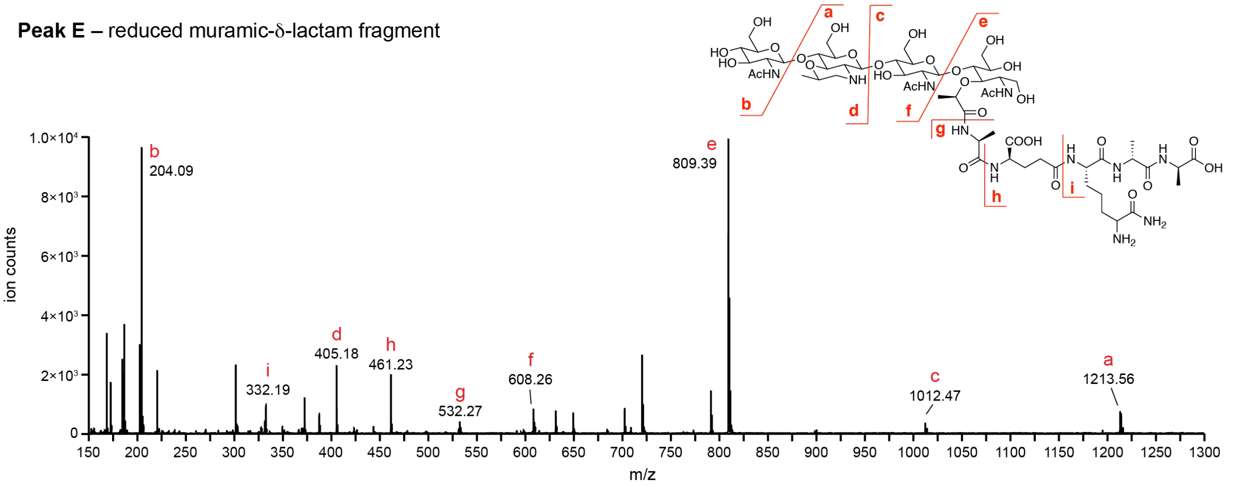


**Figure S2 (continued).** Targeted MS/MS of the peptidoglycan products formed by CwlD and PdaA. *B. subtilis* Lipid II was polymerized with SgtB and the resulting polymer treated with *Bs*CwlD and/or *Bs*PdaA as described in the Methods above. Products A, B, C, D, and E are defined as in the main text.

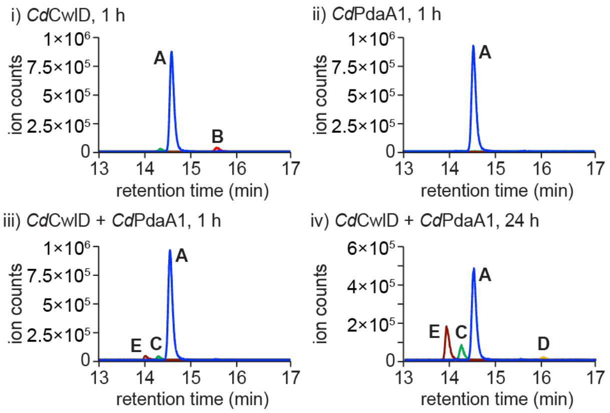


**Figure S3.** Reconstitution of *C. difficile* CwlD and PdaA1. Lipid II extracted from *B. subtilis* was polymerized with SgtB and the linear polymer treated with *Cd*CwlD and/or *Cd*PdaA1 for the indicated time. Peptidoglycan products were digested with mutanolysin and analyzed by LC-MS. The data is representative of three independent experiments.

We note that *Cd*CwlD exhibits low activity due to the absence of the activating protein GerS, which is also required for muramic-δ-lactam synthesis in *C. difficile*.[6, 7] *Cd*PdaA1 exhibited robust activity when provided substrates enriched in products B or C, as demonstrated in Figure 3, S4, and S8.

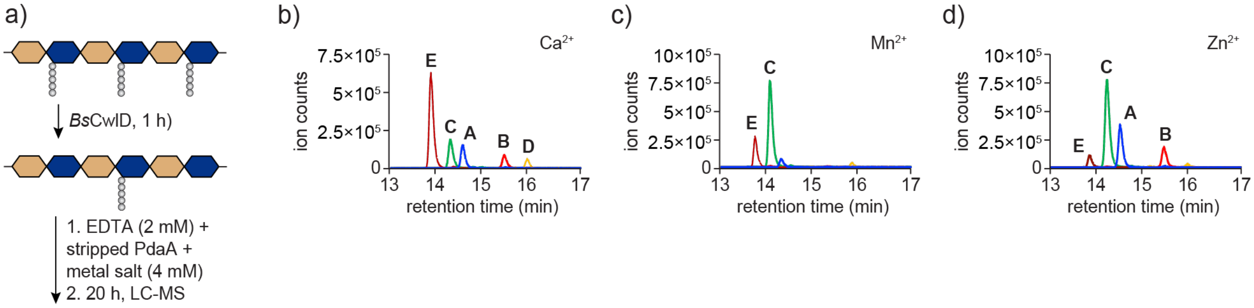


**Figure S4.** Lactam synthesis by PdaA is optimal in buffer containing $Ca^{2+}$ ions. (a) Experimental schematic. Lipid II extracted from *B. subtilis* was polymerized with SgtB and the linear polymer treated with *Bs*CwlD (2 µM) in pH 7.5 buffer without added metal ions. The reaction was quenched with 2 mM EDTA and metal-stripped *Cd*PdaA1 (2 µM) was added. Excess metal ions (4 mM, panels b,c,d) were then added to initiate the reaction and the mixture incubated at room temperature for 20 h. Peptidoglycan products were digested with mutanolysin and analyzed by LC-MS. The data is representative of two independent experiments.

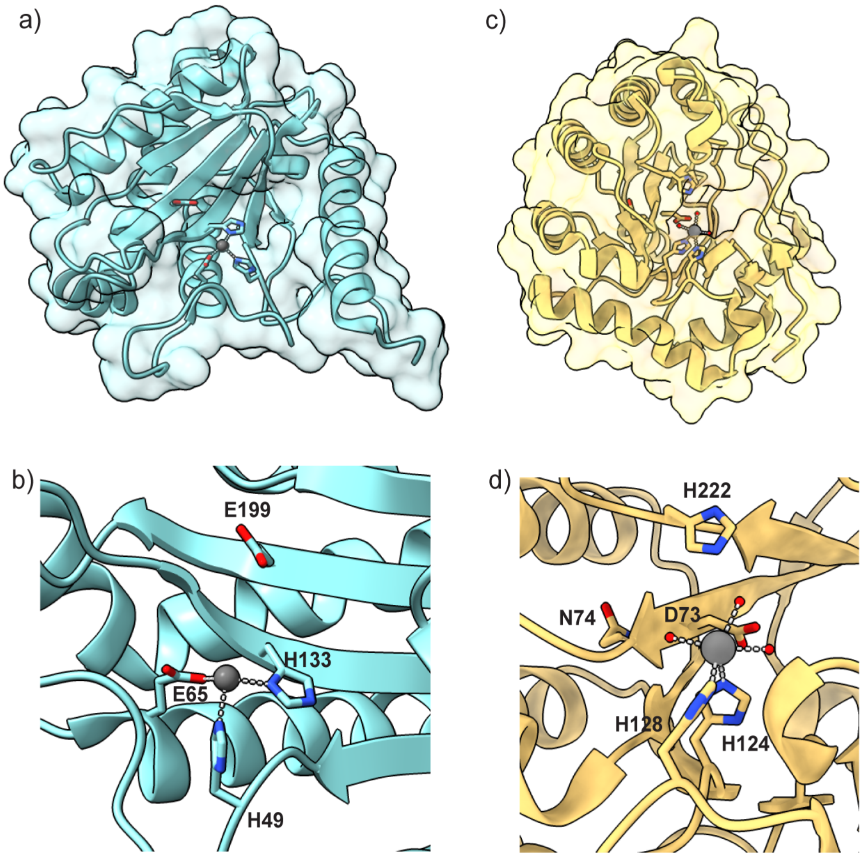


**Figure S5.** Structures of *C. difficile* CwlD[7] (panels a and b, PDB 7RAG) and *B. subtilis* PdaA[8] (panels c and d, PBD 1W1B). Conserved active site residues are indicated as sticks. The metal cofactor is shown as a gray sphere with coordinating ligands indicated with hashed lines. The red spheres in panel d indicate coordinating water molecules.

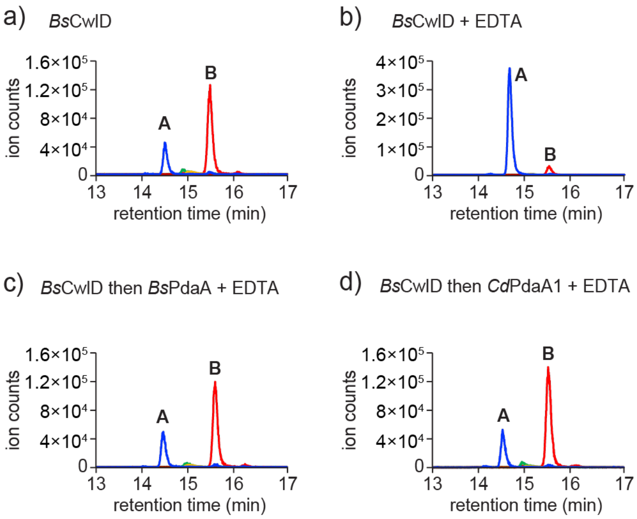


**Figure S6.** CwlD and PdaA can be quenched by addition of excess EDTA. *B. subtilis* Lipid II (20 µM) was polymerized with SgtB and the linear polymer treated with *Bs*CwlD and/or a PdaA variant (2 µM, each) for 1 h in the presence or absence of excess EDTA (10 µM). The data is representative of three independent experiments.

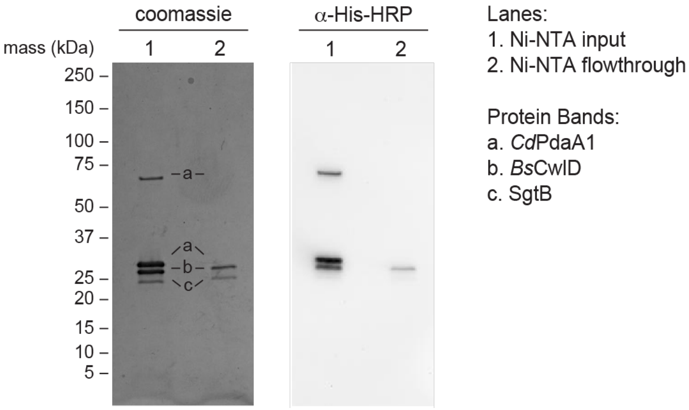


**Figure S7.** Removal of His-tagged proteins from reaction mixtures. Linear peptidoglycan was treated with *Bs*CwlD for 1 h followed by *Cd*PdaA1 for 5 min. The mixtures was passed over a plug of Ni-NTA resin and 4 µL of the eluate analyzed by SDS-PAGE with Coomassie staining or by western blot. The gel and blot shown are representative of at least three independent experiments. While a small amount of residual *Bs*CwlD remained in the sample, all *Cd*PdaA1 was removed. Similar results were obtained in reactions with *Bs*PdaA.

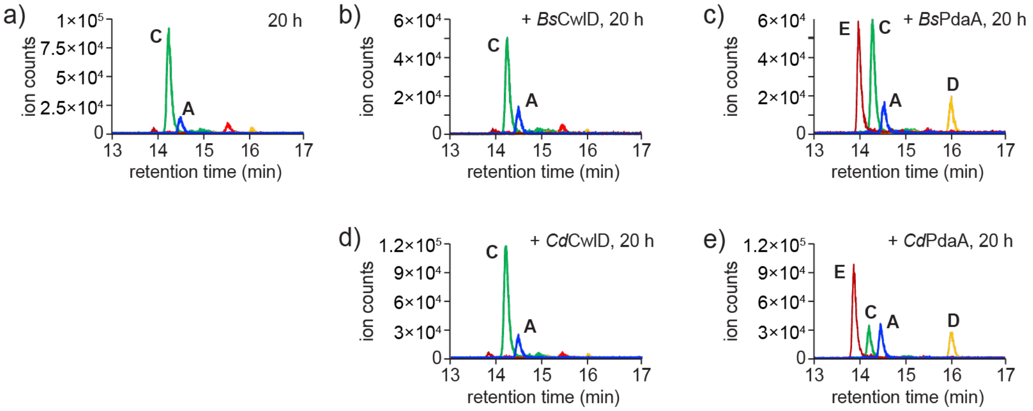


**Figure S8.** PdaA catalyzes cyclization of MurN to muramic-δ-lactam. Linear *B. subtilis* peptidoglycan was treated with *Bs*CwlD for 1 h followed by *Cd*PdaA1 for 5 min. The reaction mixture was passed over a plug of Ni-NTA to remove the proteins. The eluate was split into aliquots and analyzed by LC-MS after 20 h incubation at room temperature (a) or after 20 h treatment with the indicated enzyme (b,c,d,e). The data is representative of three independent experiments.

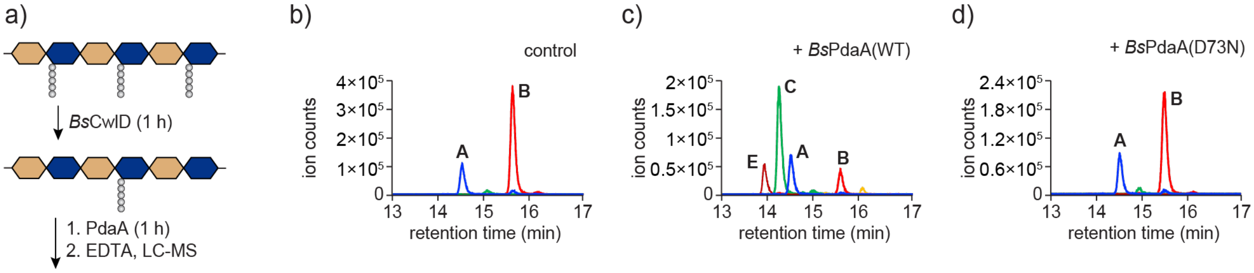


**Figure S9**. *Bs*PdaA$^{D73N}$ is an inactive deacetylase. a) Schematic of the assay used to assess PdaA deacetylase activity. Linear *B. subtilis* peptidoglycan was treated with *Bs*CwlD for 1 h followed by addition of b) no enzyme, c) wild-type *Bs*PdaA, or d) *Bs*PdaA$^{D73N}$. Reactions were incubated for 1 h at room temperature and the products analyzed by LC-MS. The data is representative of at least two independent experiments.
